# Mass spectral feature analysis of ubiquitylated peptides provides insights into probing the dark ubiquitylome

**DOI:** 10.1101/2024.05.16.594577

**Authors:** Regina M. Edgington, Damien B. Wilburn

**Author notes:** **Correspondence:** Damien B. Wilburn, 484 W 12^th^ Ave., Biological Sciences Building 880, Columbus, OH 43210.

## Abstract

Ubiquitylation is a structurally and functionally diverse post translational modification that involves the covalent attachment of the small protein ubiquitin to other protein substrates. Trypsin-based proteomics is the most common approach for globally identifying ubiquitylation sites. However, we estimate that such methods are unable to detect ~40% of ubiquitylation sites in the human proteome – i.e., “the dark ubiquitylome” – including many important for human health and disease. In this meta-analysis of three large ubiquitylomic datasets, we performed a series of bioinformatic analyses to assess experimental features that could aid in uniquely identifying site-specific ubiquitylation events. Spectral predictions from Prosit were compared to experimental spectra of tryptic ubiquitylated peptides, revealing previously uncharacterized fragmentation of the diGly scar. Analysis of the LysC-derived ubiquitylated peptides reveal systematic, multidimensional peptide fragmentation, including diagnostic b-ions from fragmentation of the LysC ubiquitin scar. Comprehensively, these findings provide diagnostic spectral signatures of modification events that could applied to new analysis methods for non-tryptic ubiquitylomics.

## Introduction

Ubiquitylation is a versatile post translational modification (PTM) where the small protein ubiquitin (Ub) is covalently attached to protein substrates. Ub is an extremely stable and hyper conserved protein, with 73 of 76 residues shared between humans and baker’s yeast.^1^ Ubiquitylation is most often associated with its role in marking proteins for degradation by the proteosome,^2^ but also participates in a broad array of cellular functions that include signaling cascades,^3^ selective autophagy,^4^ endocytosis,^5^ DNA repair,^6^ cell cycle progression,^7^ and apoptosis^8^. Consequently, improper ubiquitylation can lead to a wide breadth of cellular dysfunction and disease.^7,9^

The functional plasticity of ubiquitylation is partly due the many ways it can be attached to cognate proteins to alter their structural properties and affinity to other cellular components. In contrast to other prevalent PTMs like phosphorylation, ubiquitylation is a physically large modification (8.5 kDa) that exhibits structural diversity and widespread occurrence across the human proteome. Ub can be covalently attached to the side chains of lysine, cysteine, serine, and threonine residues, protein N-termini, and even other macromolecules such as sugars.^10–13^ After addition of a single ubiquitin to a protein reactive group (mono-ubiquitylation), further modification is possible through additional ubiquitylation events on other sites within the protein (multi-mono-ubiquitylation) or to the original ubiquitin itself (chain formation). Ubiquitin chains can be linear or branched, and the site of modification within the chain (often K6, K48, or K63) produces different functional consequences (Fig 1).^10^ The enormous structural diversity in ubiquitylation patterns contributes to its functional versatility yet makes it a challenging modification to characterize.

**Figure 1.**
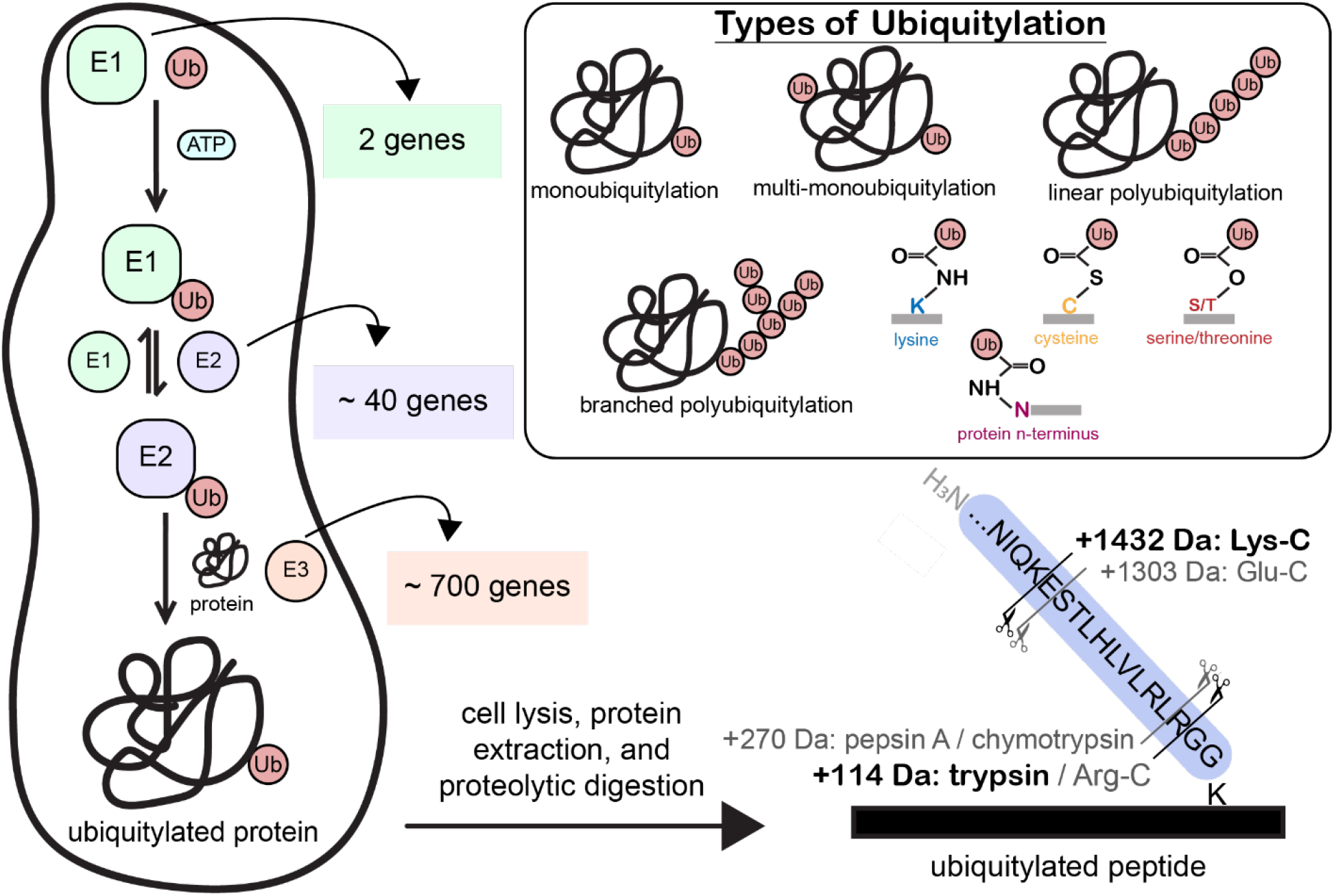
Outline of the cellular ubiquitylation mechanism and ubiquitylomics sample preparation. Ubiquitylation is a heterogeneous PTM that endogenously presents in various forms (top right). It occurs in a three-step enzymatic mechanism (shown at left). The ubiquitin-activating enzyme (E1) is charged with ubiquitin in an ATP-dependent reaction. The activated E1 enzyme transfers ubiquitin to the ubiquitin-conjugating enzyme (E2). Finally, a ubiquitin ligase enzyme (E3) facilitates the transfer of ubiquitin from the E2 enzyme to the final ubiquitylated target protein. Ubiquitylation sites can be identified using mass spectrometry. Proteins are isolated and digested with a protease enzyme, resulting in ubiquitylated peptides with a protease specific Ub scar. The minimum Ub scar from common protease enzymes is shown (bottom right).

Ubiquitylation follows a three-step enzymatic process, with each subsequent reaction showing higher specificity (**Fig 1**). In the first step, a ubiquitin-activating enzyme (E1) performs an ATP-dependent “charging” reaction whereby the C-terminus of ubiquitin is attached to the E1 catalytic cysteine by a thioester linkage. The human genome has two E1 genes that initiate all ubiquitylation events. In the second reaction, ubiquitin is transferred from the E1 to a ubiquitin-conjugating enzyme (E2) which also possesses a catalytic cysteine and uses a thioester linkage. The E2 dictates the reaction chemistry of the final ubiquitin transfer (i.e., to Lys, Ser, N-terminus, etc.), and ~40 human E2 genes perform these different modification types. Finally, a ubiquitin ligase enzyme (E3) coordinates the transfer of Ub from the E2 to the final target protein by either acting as scaffold for direct transfer (e.g. RING-type E3s) or forming an E3~Ub intermediate before transferring Ub to the substrate(e.g. HECT and RBR-type E3s).^10^ Using one of the ~700 human E3 genes, the structure of the E2-E3-substrate ternary complex ultimately determines where the modification is added, and this large combinatorial space creates tremendous opportunity for much of the proteome to potentially be ubiquitylated with site-specificity.

Historically, ubiquitylation of specific proteins has been assessed through western blotting^11^, various enrichment strategies (immunoprecipitation, ubiquitin binding domains, etc. )^14^, and/or targeted *in vitro* ubiquitylation reactions. These experiments offer valuable insights into modifications on the protein level, but usually cannot provide site localization information without further experimental validation (e.g. site-directed mutagenesis). Liquid chromatography paired with tandem mass spectrometry (LC-MS/MS) is the “gold standard” approach for PTM localization, with advances in instrumentation and data analytics enabling “proteome-wide” PTM detection. The typical workflow of such experiments involves digesting protein samples with a proteolytic enzyme, enriching for modified peptides using modification-specific antibodies, and performing LC-MS/MS analysis.^15–17^ Peptide fragmentation patterns can be used for PTM site localization, offering amino acid level resolution of modification events.

Proteolysis of ubiquitylated proteins results in crosslinked peptides that include a C-terminal scar from Ub at the modification site, and digestion with trypsin leaves only a small di-glycine (diGly) peptide at the ubiquitylated residue (+114 Da; **Fig 1**).^18^ While the diGly modification is relatively easy to observe by LC/MS-MS, it has several limitations for detecting specific ubiquitylation sites. First, Ub-like proteins NEDD8 and ISG15 also produce a diGly scar when modified proteins are trypsinized, and therefore it cannot be used to unambiguously distinguish between these similar PTMs (which are known to occur on the same proteins).^19^ Additionally, commercially available antibodies are only available for diGly-modified lysine and are reported to have variable affinity depending on the cognate peptide sequence^12,19,20^ Most importantly, detected site-specific modifications are inherently biased towards regions of the proteome that produce tryptic peptides well suited for LC/MS-MS analysis. Because trypsin cleaves peptide bonds on the C-terminal side of arginine and unmodified lysine residues, peptides produced from basic rich regions of the proteome are too small to be unambiguously assigned to a precursor protein, while limitations in instrumentation make it challenging to identify long peptides from basic depleted regions. It is estimated that tryptic peptides can only provide ~60% sequence coverage of the unmodified human proteome, posing challenges in PTM studies where high coverage of diagnostic fragment ions is critical for confident modification assignment.^21^ Consequently, we argue that new experimental and computational methods are needed for both detection and quantification of the dark ubiquitylome: the many site-specific ubiquitylation events that are inaccessible to trypsin-based proteomic methods .

We address prevailing challenges of characterizing the dark ubiquitylome by conducting a meta-analysis of three recently published ubiquitylomics datasets. First, search results from these studies were aggregated and used to estimate the size of the dark ubiquitylome. Evaluating data produced by standard methods, spectral features of diGly peptides were systematically evaluated in comparison to their unmodified counterparts and non-trivial peptide fragmentation of the scar itself was identified. To assess the utility of alternative proteases, MS2 spectra of ubiquitylated peptides with a LysC Ub scar were analyzed, providing insight into the multidimensional but systematic fragmentation pattern of these crosslinked peptides. We propose that future integration of these features into existing search engine and deep learning platforms may provide a roadmap for more comprehensive ubiquitylomics analysis.

## Methods

### Datasets

Raw mass spectrometry data was aggregated from three ubiquitylomics studies. Data was downloaded from Steger *et al*.^15^ from the PRIDE partner repository with the dataset identifier PXD023889. In this study, proteins were isolated from three cell lines (HTC116, MM.1S, and Jurkat 6.1) followed by digestion with a trypsin/Lys-C mixture. HTC116 cell lysates were subject to high-pH reversed-phase offline fractionation. DiGly peptides were enriched using a KGG antibody-bead conjugate, which targets the lysine-G-G (diGly) epitope.

LC-MS/MS analyses were performed on an EASY-nLC 1200 system (ThermoFisher) coupled with a Q Exactive HF-X mass spectrometer (ThermoFisher) using DDA and DIA methods.

Data from Hansen *et al*.^16^ was downloaded from the dataset identifier PXD019854. Cell lyates from HEK293 and U2OS cells were digested with trypsin. DiGly peptides were enriched using a KGG antibody-bead conjugate and high-pH reversed-phase offline fractionation was performed. LC-MS/MS was conducted employing an identical instrumental setup and comparable data acquisition methodologies as described by Steger *et al*.^15^

Ubiquitylomics data from Akimov *et al*.^17^ was downloaded from the dataset identifier PDX006201. The authors treated multiple human cell lines (Hep2, Jurkat, HEK-293T) with one of a set of proteome inhibitors (DMSO control, bortezomib, b-AP15). Proteins were first digested with LysC, and LysC-derived ubiquitylated peptides were enriched using the UbiSite monoclonal antibody that was raised against the 13 amino acid C-terminal Ub scar peptide. Following enrichment, an aliquot of peptides were further digested with trypsin. High-pH reversed-phase offline fractionation was performed for both LysC and tryptic peptides, followed by analysis using an EASY-nLC 1000 system (ThermoFisher) connected to a Q Exactive HF mass spectrometer (ThermoFisher) using DDA methods. Tryptic peptides were further analyzed using DIA methods.

Spectral libraries of tryptic ubiquitylated peptides from Steger *at al*.,^15^ Hansen *et al*.,^16^ and Akimov *et al*.^17^ were searched using a common data analysis pipeline. Aggregated search results were compared to the theoretical human ubiquitylome. DDA data from the Hep2 cells treated with bortezomib from Akimov *et al*.^17^ were used for the spectral analyses performed in this study.

### Search and Validation

Raw files were converted to mzML using ThermoRawFileParser.^22^ Searches and validation were performed in Fragpipe, version 20.0. For searching, MSFragger (v 3.8) was used with deisotoping and deneutral loss features enabled.^23^ Data were searched against the Uniprot human reference proteome (downloaded 06/12/2023) with decoys and common contaminants added using Philosopher.^24^ Searches were performed with intact N-terminal methionine residues. Protease cleavage (trypsin or Lys-C) was assigned based on protease specificity, with no digestion before proline residues. Peptides between 6-50 residues with m/z values between 500-5000 considered. Up to three missed cleavages were permitted per peptide. Mass tolerance values of 50 ppm and 20 ppm were used for MS1 and MS2 measurements respectively. Cysteine carbamidomethylation was set as a fixed modification, and the following variable modifications were considered: methionine oxidation, protein N-terminal acetylation, glutamine ammonia loss, and glutamic acid water loss. For tryptic and LysC peptides, respectively, either the Ub two amino acid scar (+114 Da) or thirteen amino acid scar (+1432 Da) was included as a variable modification. Up to three variable modification events were permitted per peptide. PSM validation was performed using MSBooster^25^ and Percolator^26^. PTMProphet was enabled for modification site localization.^27^ Data was filtered to a 1% peptide-spectrum match (PSM) level false discovery rate. Search results were linked to experimental spectra using BiblioSpec (v 2.0) from Skyline.^28^ Encyclopedia^29^ (v 2.12.30) was used to generate experimental DLIB libraries that were imported into Jupyter notebooks for analysis.

### Custom Bioinformatic Analysis

Custom scripts in Python were used for bioinformatic and spectral feature analysis. Detection probability by mass was estimated by cubic spline fitting using the scipy function interp1d. Experimental spectra were compared to theoretical spectra using the cosine similarity function (equation 1):

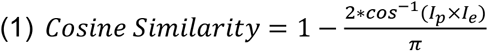

In this equation, I_p_ represents predicted ion intensities and I_e_ represents experimental ion intensities. Similarity scores were computed with a 20 ppm mass tolerance. Spectra were L2-normalized prior to computing cosine similarity.

Prosit predicted libraries were generated using Prosit 2020 intensity HCD and Prosit 2019 iRT models.^30^ Prosit collision energies were optimized by predicting libraries of experimentally observed precursors at variable collision energies (25-35 for the tryptic peptides and 25-42 for the LysC peptides). Spectral comparison scores between predicted and experimental spectra of unmodified peptides were computed, with the average highest scoring collision used in subsequent analyses: NCE 34 was for tryptic predictions and NCE 37 for LysC predictions. Mass shifts were applied to fragment ions as necessary for alignment of fragmentation ladders between ubiquitylated and unmodified peptides.

To analyze the fragmentation of the tryptic Ub peptides, Prosit predictions of unmodified and diGly peptides were directly compared to experimental spectra. Cosine similarity scores were computed based on the location of the diGly modification within the peptide. The theoretical m/z values of modified fragment ions were systematically varied based on chemically feasible mass loss events. Using these values, cosine similarity scores were computed between experimental and theoretical modified fragment ions.

The LysC Ub peptide fragment ions were annotated by considering peptide fragmentation along the main peptide backbone and along the 13 amino acid Ub scar. Fragment ion m/z values corresponding to b/y-ion ladders for the peptides were calculated. The Ub scar length was varied from 0-13 amino acids on all fragment ions with the Ub modification. Fragment ion charge states up to +3 were considered. Theoretical fragment ions were matched to experimental fragment ions within a 20-ppm mass tolerance. Fragmentation of the experimental LysC crosslinked peptides were compared to the Prosit predicted spectra of the LysC Ub scar (peptide ESTLHLVLRLRGG +2H).

## Results and Discussion

### Bioinformatic interrogation of the dark ubiquitylome

To estimate the undetectable fraction of the human ubiquitylome using trypsin-based methods – i.e., the dark ubiquitylome – we leveraged available data from three large ubiquitylomics studies. In Akimov *et al*.,^17^ proteome samples were first proteolyzed with LysC to produce peptides with longer Ub scars against which a monoclonal antibody was raised (Ubisite); following enrichment, peptides were further digested with trypsin to yield diGly peptides. In both Hansen *et al*.^16^ and Steger *et al*.^15^, offline fractionated DDA libraries of diGly-enriched peptides were generated for quantitative DIA analysis of ubiquitylome perturbation. Raw data from the three studies was searched using a common informatics pipeline (see methods). We identified a total of 120,567 Ub-modified lysines across the human proteome on peptides without miscleavages (except at the modified lysine), or ~19% of the theoretical max (~640K). As expected, the detection rate of ubiquitylated peptides was highly dependent on the mass of the modified peptide, ranging between 0% and 31.3%. If we assume that the maximum detection rate is close to the true proteome-wide average, then the expected ubiquitylome would be closer to 200K peptides and suggest that ~40% of site-specific ubiquitylation events are undetectable by trypsin-based methods (**Fig 2A**). While this value is likely an overestimate given that the cell lines used by the authors do not express all substrates, E2s, or E3s, it is striking that this value is nearly identical to reported calculations for the unmodified dark proteome.^21^

**Figure 2.**
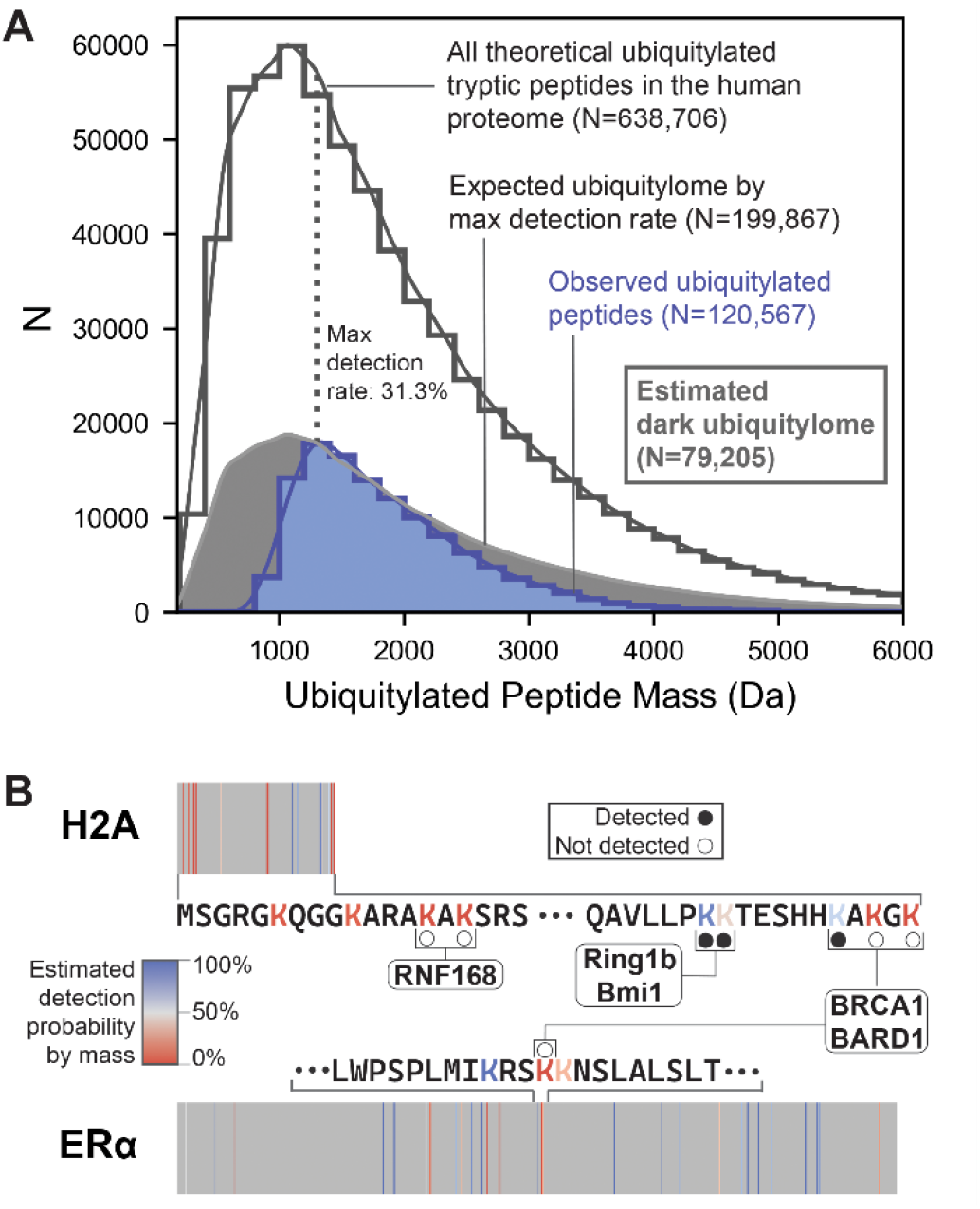
Bioinformatic characterization of the dark ubiquitylome. (A) There exist >600K lysine residues in the human proteome that could be potentially ubiquitylated, and by *in silico* digestion, we calculated the expected mass distribution of all theoretical ubiquitylated peptides in the human proteome (without missed cleavages). Compared to the ~120K observed sites in available ubiquitylomics datasets, we estimated that current methods maximally detect ~31.2% of ubiquitylated peptides within any mass bin, and ~79K or 40% of ubiquitylation sites are undetectable with trypsin-based proteomics. (B) Many important site-specific ubiquitylation events for human health and disease are found within the dark ubiquitylome, including some of the most biochemically well-characterized examples such as BRCA1/BARD1 ubiquitylation of histone H2A and estrogen receptor α.

While the most well-known functions of Ub are chain formation and protein degradation, mono-Ub is a far more abundant modification and critical for proper cellular function. For example, ~5-15% of histone H2A is mono-ubiquitylated in human cells and present at concentrations ~20-80% of free Ub.^31,32^ At least three different E3 enzymes specifically mono-ubiquitylate different clusters of lysine residues on the disordered tails of H2A, yet only 3 out of the 7 verified modification sites were detected from the aggregated ubiquitylomics data (**Fig 2B**). Most mono-Ub is added to H2A K118/K119 by Ring1b/BMI1 for transcriptional silencing. By contrast, BRCA1/BARD1 are well-known tumor suppressors associated with breast cancer that specifically ubiquitylate one of the nearby K125/K127/K129 residues to recruit DNA repair machinery.^33^ RNF168 also contributes to DNA repair, but through ubiquitylation of K13/K15 and a likely different mechanism.^34^ Mono-ubiquitylation of other histone tails also play diverse and important roles in chromatin regulation.^35^ As another class of DNA binding protein, transcription factors also commonly possess basic regions whose functionality is modulated by mono-Ub, such as K302 on estrogen receptor α that is also ubiquitylated by BRCA1/BARD1 (Fig 2B).^36^

Precise site-localization of a PTM to a specific residue on a peptide can be challenging if there are multiple potential attachment sites and relatively few diagnostic fragment ions in the MS2 spectrum.

Quantitative interrogation of these difficult-to-probe regions in the ubiquitylome requires innovation in sample preparation, LC/MS-MS analysis, and integration of additional experimental features into search engines. To begin addressing this challenge, we performed a series of exploratory data analyses to identify how ubiquitylated peptides with different scar lengths vary compared to each and unmodified counterparts predicted with Prosit.

### Analysis of diGly-modified tryptic peptides

The spectral features of ubiquitylated tryptic peptides with a diGly amino acid Ub scar were investigated. We utilized data from Hep2 cells treated with bortezomib, as reported by Akimov *et al*.,^17^ for C18 retention time (RT) and fragmentation feature analysis. In this experiment, cell lysates were digested with LysC, and Ub peptides were enriched with the UbiSite antibody. After enrichment, LysC peptides were digested with trypsin, resulting in a mixture of diGly peptides (~31K) and unmodified peptides (~70K) released from internal arginine cleavages. From this dataset, we identified 1569 peptide sequences where measurements were recorded with and without the modification, allowing for direct assessment of the RT effects from the diGly motif. RT alignment between the two forms revealed a strong correlation with a mean and median RT shifts of 2.1 and 1.5 minutes, respectively (**Fig 3A-B**). We hypothesize that this consistent RT increase is produced by the diGly modification extending the lysine side chain and increasing interactions with the C18 resin without altering the net charge (due to the N-terminal amine of the diGly).

**Figure 3.**
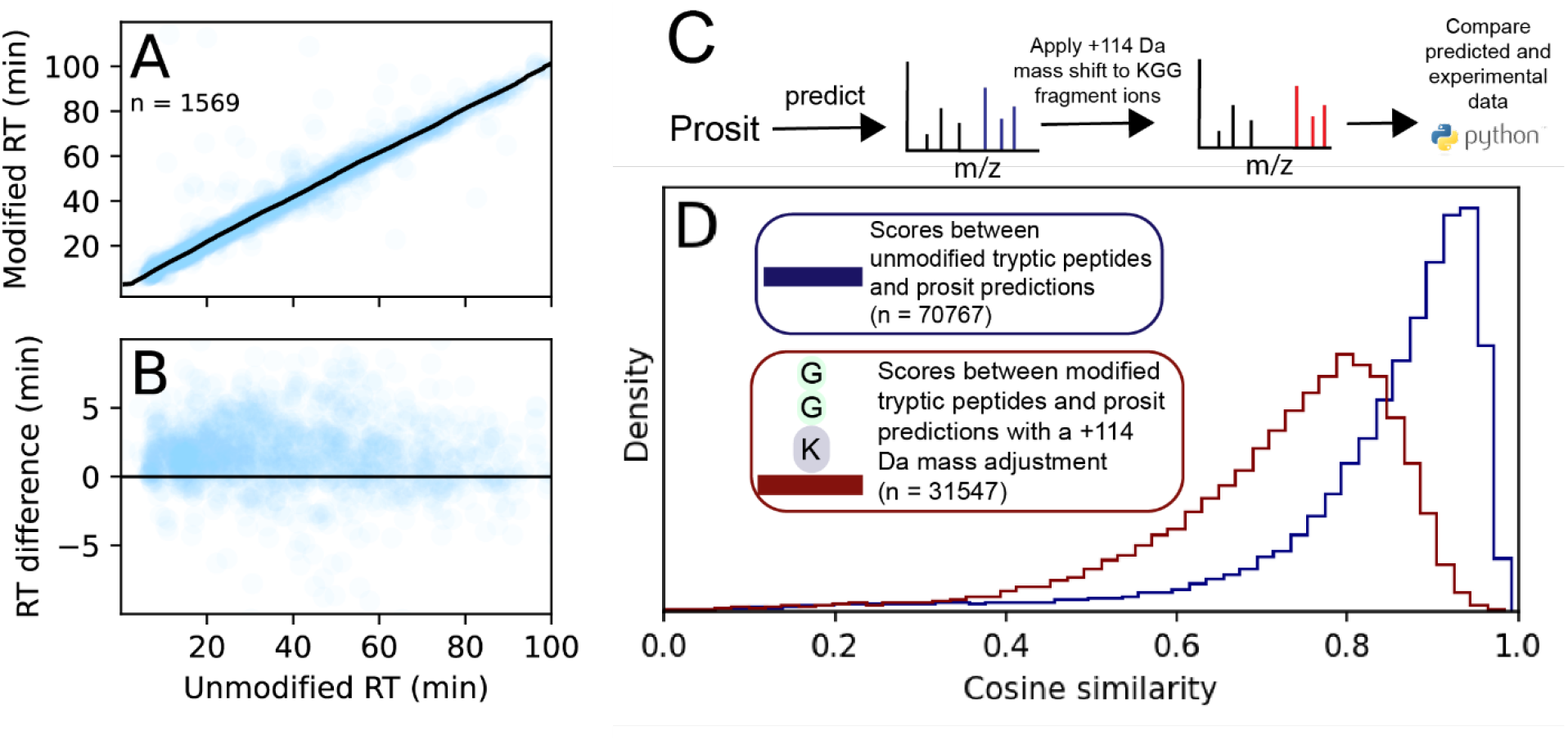
Retention time analysis and global comparison of tryptic Ub peptides. (A) KDE-based retention time alignment between the 1569 shared unmodified and diGly peptides in the Akimov *et al*. dataset. (B) RT shift between unmodified and ubiquitylated peptides, with modified peptides eluting an average of 2.1 minutes earlier than their tryptic counterpart (median RT shift = 1.5 min). (C) Workflow for predicting modified spectra with Prosit where a +114 Da mass shift fromdiGly is applied to the fragment ions that contain the Ub scar. (D) Cosine similarity distributions for all peptides in the dataset computed between Prosit and experimental spectra.

We subsequently aimed to explore how the diGly modification alters peptide fragmentation in MS2 spectra. To improve our statistics, we desired to analyze the complete set of ~31K diGly peptides, and as such used the deep learning model Prosit to generate spectral predictions for the unmodified form of peptide sequences observed in the dataset (regardless of modification state). Prosit was largely trained on unmodified tryptic peptides and can predict the fragmentation of such peptides with high accuracy.^34^ Observed and predicted b/y-ion ladders were aligned to calculate cosine similarity score distributions. We found that ubiquitylated spectra (median = 0.75) scored substantially worse than unmodified versions (median = 0.88) (**Fig 3C-D**), hinting at a potential systematic difference in fragmentation of the diGly modified peptides.

Given this effect, we next investigated if the positional location of the diGly modification might have a measurable effect on fragmentation. Disaggregating the cosine similarity distribution by peptide length and position of the diGly supported that – regardless of peptide length – there is lower similarity between modified spectra and unmodified predictions as the modification approaches the C-terminus (**Fig 4A, S1-3**). Comparison of fragment ion intensities identified that Prosit consistently overestimated the intensities of b/y-ions that contain the diGly modification (**Fig 4B-C**). Based on this result, we hypothesized that fragment ions with the Ub scar may be undergoing an extended fragmentation process after ion activation.

**Figure 4.**
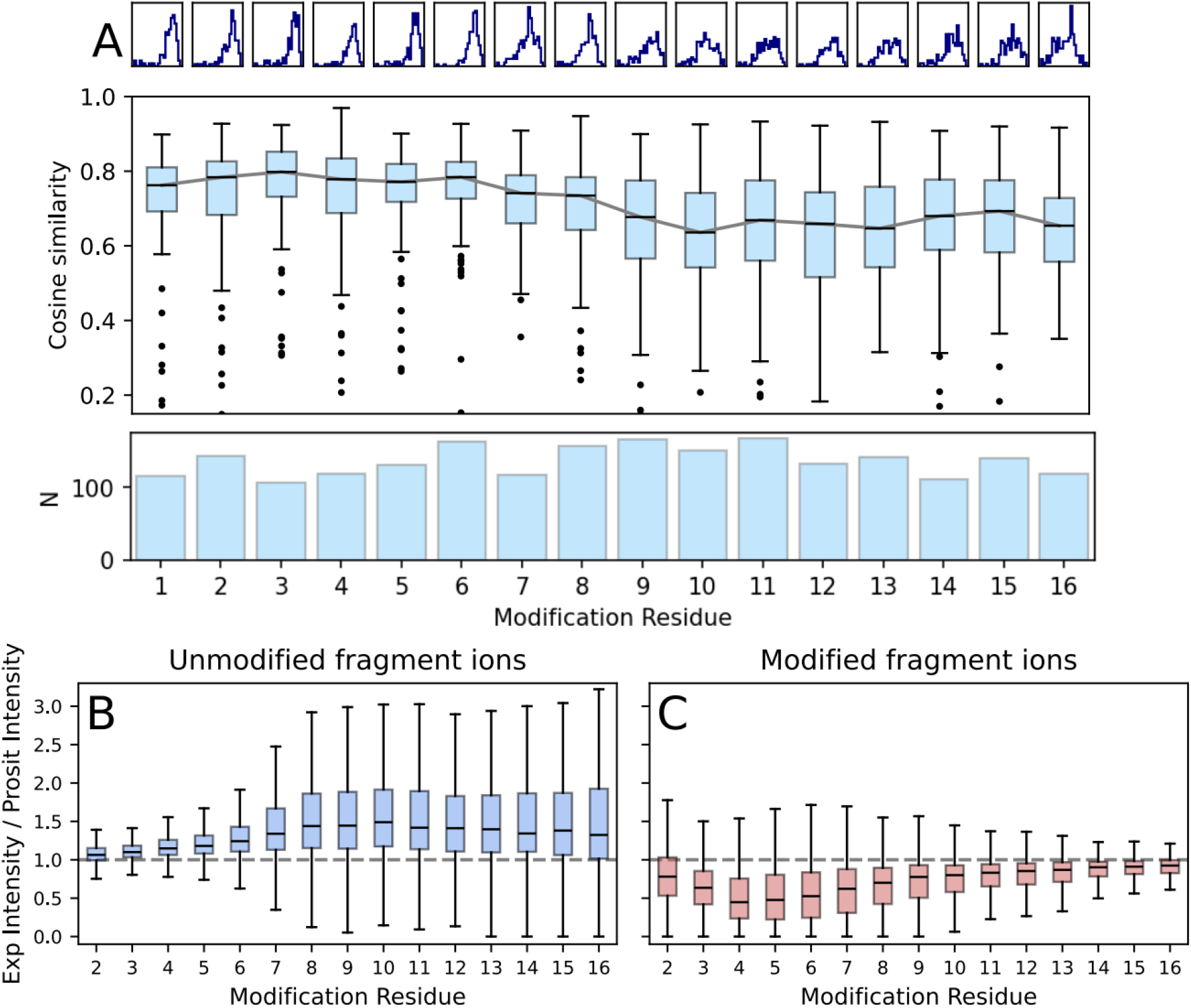
Cosine similarity score between experimental and predicted spectra is related to modification residue. (A) Cosine similarity scores for all modified peptides 17 amino acids long. Score distributions are shown at the top. Boxes represent the 25^th^ and 75^th^ percentile of scores. Outliers are ± 1.5 times the interquartile rage. The solid grey line represents median score values for each modification residue. The number of peptides scored at each residue is shown in the bottom bar graph. (B-C) Examination of the ratios of experimental to Prosit predicted ion intensities for all (B) unmodified and (C) modified fragment ions with modification residues between 2-16. Total intensities summed across each individual spectrum, and outliers are not shown.

MS2 spectra of tryptic peptides are typically dominated by y-type ions, especially when fragmented by beam-type CID (HCD).^37^ As the Ub modification gets closer to the C-terminus of the peptide, more y-ions retain the scar. We hypothesized that the C-terminally modified peptides were scoring worse because more of their spectrum is composed of fragment ions that contain the Ub scar, and these ions are undergoing an alternative degradation pathway. To test this hypothesis, the experimental spectra of C-terminally modified peptides were scored against the predicted modified fragment ions with different mass loss values (based on all possible monoisotopic compositional losses) (**Fig 5B-C**). We observed three major degradation products: –18 water loss, –57 glycine loss, and –114 diglycine loss. Water loss is commonly seen in peptide fragmentation, and it remains unclear whether this observation was related to the Ub scar or not. However, the glycine and diglycine losses resulted from fragmentation at the peptide bonds along the diGly Ub scar. Both the observed RT shift and predictable neutral loss species are potentially useful diagnostic features to help researchers increase confidence in site-specific localizations of Ub on tryptic peptides.

**Figure 5.**
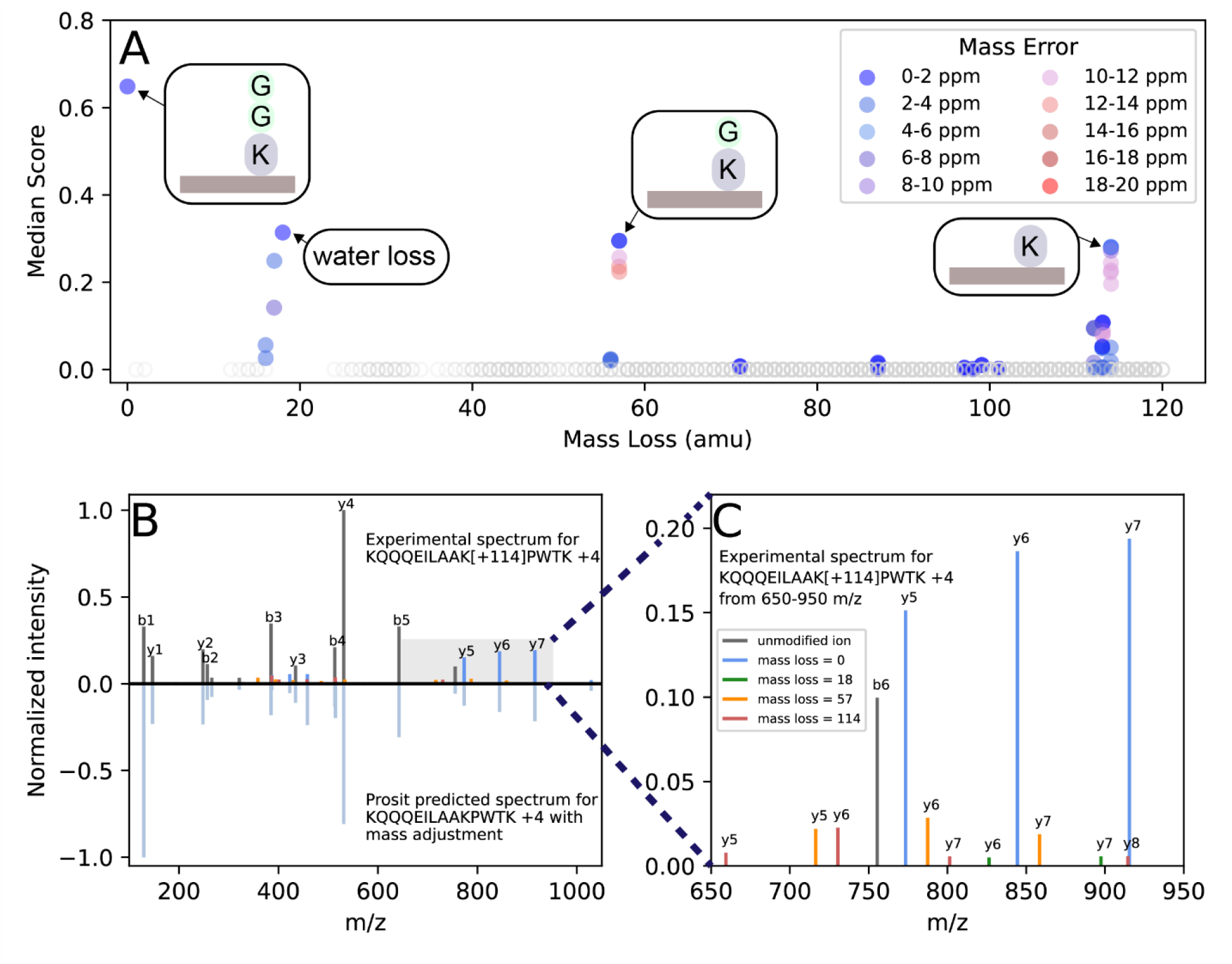
Tryptic peptides fragment along diGly Ub scar. (A) Median cosine similarity scores between experimental spectra and Prosit predicted modified fragment ion intensity at different mass loss values. Monoisotopic mass loss values were calculated from different chemically feasible combinations of carbon, hydrogen, nitrogen, and oxygen atoms. Only peptides with a C-terminal modification are scored (modification residue ≥ 10). Open grey circles indicate a median score of 0. Median ppm error values are shown for nonzero score values. (B) Mirror plot of experimental spectrum and mass-adjusted Prosit-predicted spectrum of the modified peptide KQQQEILAAK[+114]PWTK +4H. (C) Increased magnification of the 650-960 m/z region from B with scar fragmentation colored.

### LysC Ub Peptides

The proximity of protease cleavage sites to any residue position in the proteome is an unalterable biological constraint, and alternative proteases may be useful in observing ubiquitylation events currently inaccessible by trypsin. However, as substantial scar fragmentation was observed on a simple diglycine, use of any other protease will increase the Ub scar length (**Fig 1**) and we anticipate will produce even more complex fragment ion spectra. To assess fragmentation patterns of ubiquitylated peptides with longer Ub scars, spectra of LysC peptides with a 13 amino acid Ub scar were systematically evaluated using data from the Hep2 cells treated with bortezomib from Akimov *et al*. When compared to the matched tryptic dataset, we identified 1940 peptide sequences that were detected with both lengths of scars. Of these peptides, nearly 90% of spectra exhibited an increased charge state of either +1 or +2 relative to their tryptic counterparts. The elevated charge state in LysC-derived peptides was expected given the highly basic nature of the LysC Ub scar, which contains two arginine residues and one histidine residue.

Like most crosslinked peptides, the structure of LysC-derived Ub peptides allows for multidimensional fragmentation. “Traditional” peptide fragmentation occurs along the backbone of the cognate peptide, but simultaneously fragmentation can also occur along the 13 residue Ub scar. To examine these complex spectra, we developed scripts to classify observed fragment ions as one of four types: fragment ions that do not contain the Ub modification, fragment ions that contain the intact 13 amino acid Ub scar, fragment ions that contain a fragmented Ub scar (< 13 amino acids long), and b-ions resulting from fragmentation of the Ub scar (**Fig 6A**). Of fragment ions that contained the modification site, 48.1% were b-ions and 51.9% are y-ions. As expected, the fragment ions with the scar typically had an elevated charge state: 39.3% were +2 and 53.5% were +3. Across all fragment ions that contained the modification site, 28% of ion intensity were from fragments with an intact scar compared to 72% with a fragmented scar. We also used Prosit to predict the spectrum of the Ub scar (peptide ESTLHLVLRLRGG +2H), and to our surprise, many b-ions from this scar were detected across LysC spectra (**Fig 6B-C**). As larger m/z fragment ions are less susceptible to interference, we identified 7 diagnostic b-ions that we observed were reproducibly detected between LysC-derived spectra (**Fig 6D**).

**Figure 6.**
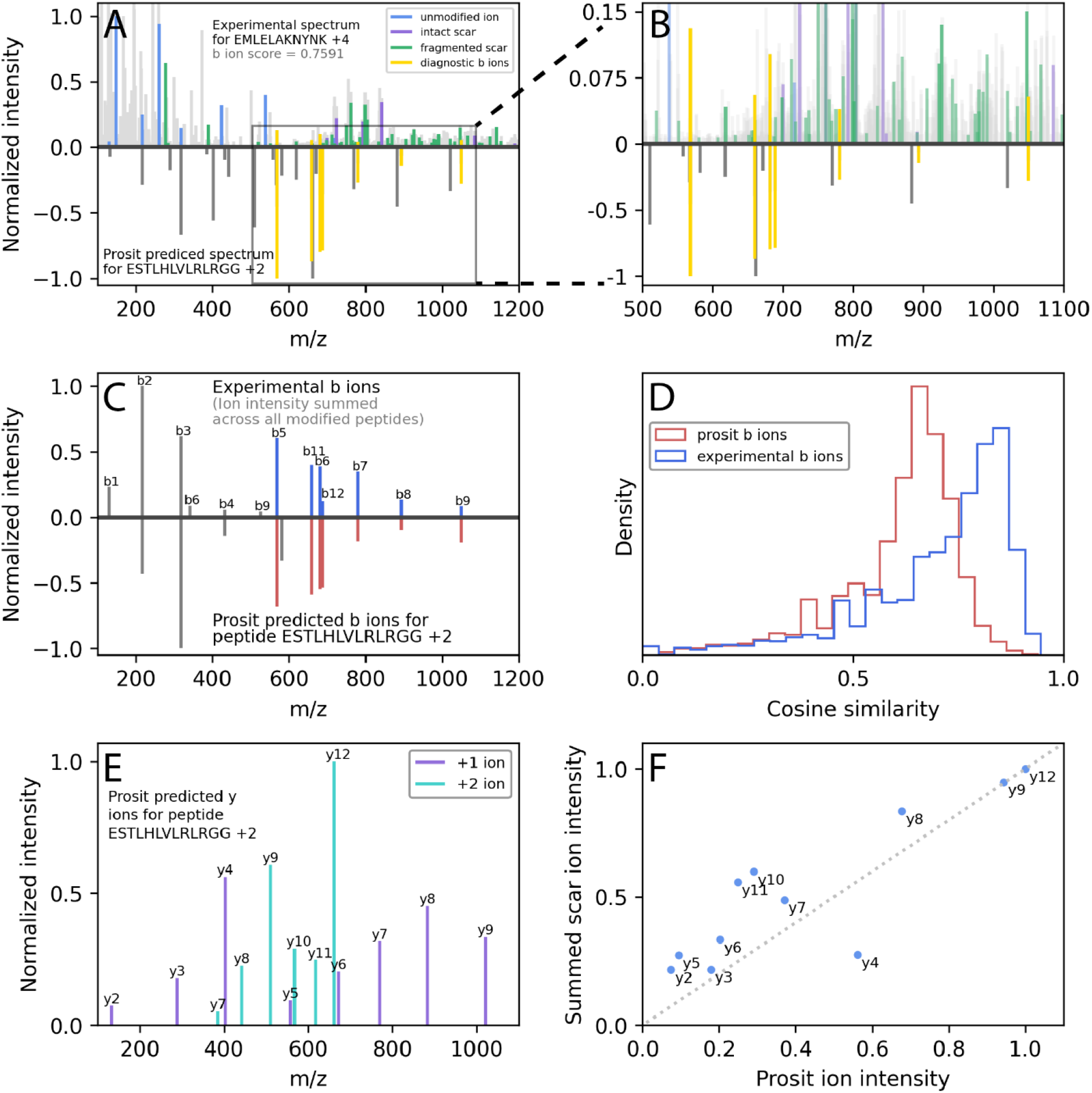
Crosslinked LysC peptides show correlative fragmentation in two dimensions. (A) Experimental spectrum of ubiquitylated LysC peptide EMLELAK[+1432]NYNK +4H with annotated fragment ions (top). Blue ions are peptide fragments that do not contain the modification site; purple ions are fragments that contain the intact 13 amino acid Ub scar; green ions are fragments that contain a fragmented Ub scar (< 13 amino acids); gold ions are diagnostic b-ions resulting from fragmentation of the LysC Ub scar. The bottom mirror plot spectra shows the Prosit prediction of the LysC Ub scar (peptide ESTLHLVLRLRGG +2H), with (B) a zoom of the 500-1100 m/z range with emphasis on the diagnostic b-ions. (C). Scar b ion intensities summed across all modified spectra (top) and predicted by Prosit (bottom), with diagnostic b-ions colored in blue/red. (D) Cosine similarity score distributions between experimental modified spectra and diagnostic b ion intensities from C. (E) Prosit predicted y ions of the LysC Ub scar (peptide ESTLHLVLRLRGG +2H). (F) Correlation between the Prosit y-ion intensities and experimental intensities of fragment ions that contain a fragmented Ub scar. Experimental ions were categorized based on the scar attachment after fragmentation.

While b-ions from the Ub scar are freely released into the gas phase, the corresponding y-ions remain attached to the cognate peptide and are not as simply analyzed. However, we still desired to investigate if these signals also offer useful insights for strategies to improve statistical confidence of Ub site localization.

We compared Prosit predictions of Ub scar y-ion intensities to summed experimental fragment ions that have a fragmented Ub scar (i.e. experimental fragment ions were categorized by which scar y-ion was retained after activation). The collective observed fragmentation of the Ub scar was found to be related to the predicted fragmentation of the scar by itself (**Figs 6E-F, S4-5**). This result supports that fragmentation of peptides with a complex Ub scar is complex, but systematic.

Importantly, these findings suggest that fragmentation patterns inferred on naïve peptides are potentially transferrable to crosslinked cognates. In addition to important natural protein crosslinks such as Ub and Ub-like proteins, *in vitro* crosslinking is an invaluable experimental tool to identify transient protein-protein interactions under natural physiological contexts. However, crosslinking proteomic data are complex and researchers often incorporate higher orders of analytical resolution (e.g. MS3).^38^ Deep learning models typically require enormously large datasets to train, and the exceptional performance of Prosit partly due to the availability of high-quality data from synthetic peptides generated by the ProteomeTools Project.^30,39,40^ It would be challenging to generate an equivalent synthetic dataset for crosslinked peptides, but our findings on Ub suggest that transfer learning or other iterative machine learning approaches may be sufficient to extend existing architectures to this important application.

## Conclusions

Ubiquitylation is a structurally and functionally diverse PTM, yet we estimate that current trypsin-based proteomic methods are unable to identify ~40% of the human ubiquitylome. To expand our knowledge on the diverse impact of this modification, alternative analytical techniques are needed for identification and quantification. Limitations in trypsin proteolysis suggest that non-tryptic proteases could help capture modification events that are currently inaccessible, but non-tryptic digestion produces crosslinked peptides with longer Ub scars. Although the MS2 spectra of these peptides are complicated, our analyses suggest that they are surprisingly interpretable. In the future, we believe deep learning algorithms could be leveraged to directly predict the spectra of these peptides, which has possible applications for new database search algorithms, studying protein-protein interactions, and – most critically – illuminating the dark ubiquitylome.

## Acknowledgements

This work is supported by funding from the National Institutes of Health (R00-HD090201 and R35-GM150583 to D.B.W.) and an OSU ERIK JobsOHIO research award. We thank Dr. Rachel Klevit for her guidance in the preparation of this manuscript. We also extend thanks to James Alltop, Trenton Winters, and other members of the Wilburn lab for their assistance in the writing process.

