## Supplemental Figures for "Mass spectral feature analysis of ubiquitylated peptides provides insights into probing the dark ubiquitylome"

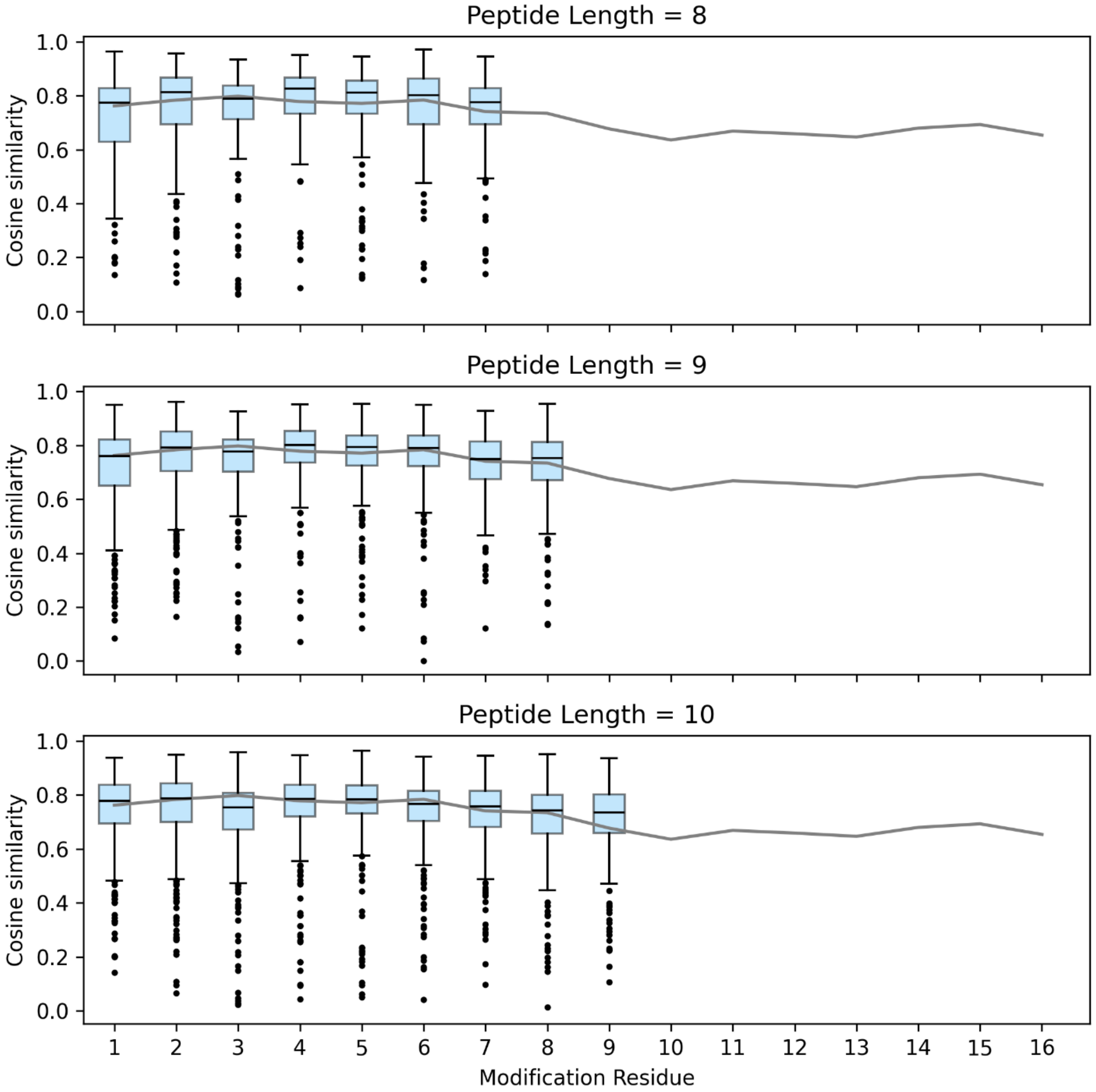


**Figure S1.** Cosine similarity scores for all modified peptides 8, 9, and 10 amino acids long. Boxes represent the 25th and 75th percentile of scores. Outliers are ± 1.5 times the interquartile rage. The solid grey line represents median score values for each modification residue for all modified peptides 17 amino acids long categorized by modification residue.

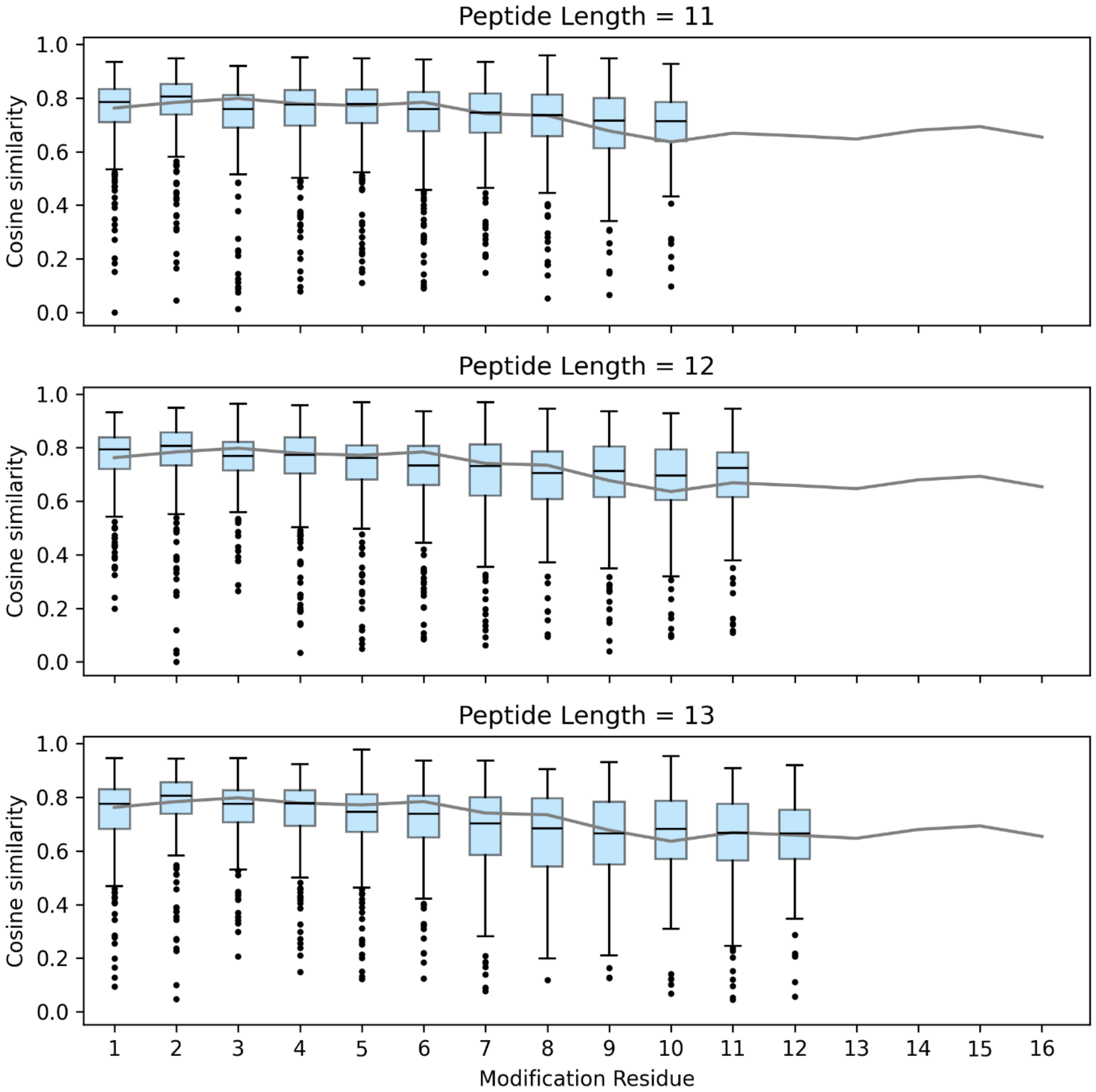


**Figure S2.** Cosine similarity scores for all modified peptides 11, 12, and 13 amino acids long. The solid grey line represents median score values for each modification residue for all modified peptides 17 amino acids long categorized by modification residue.

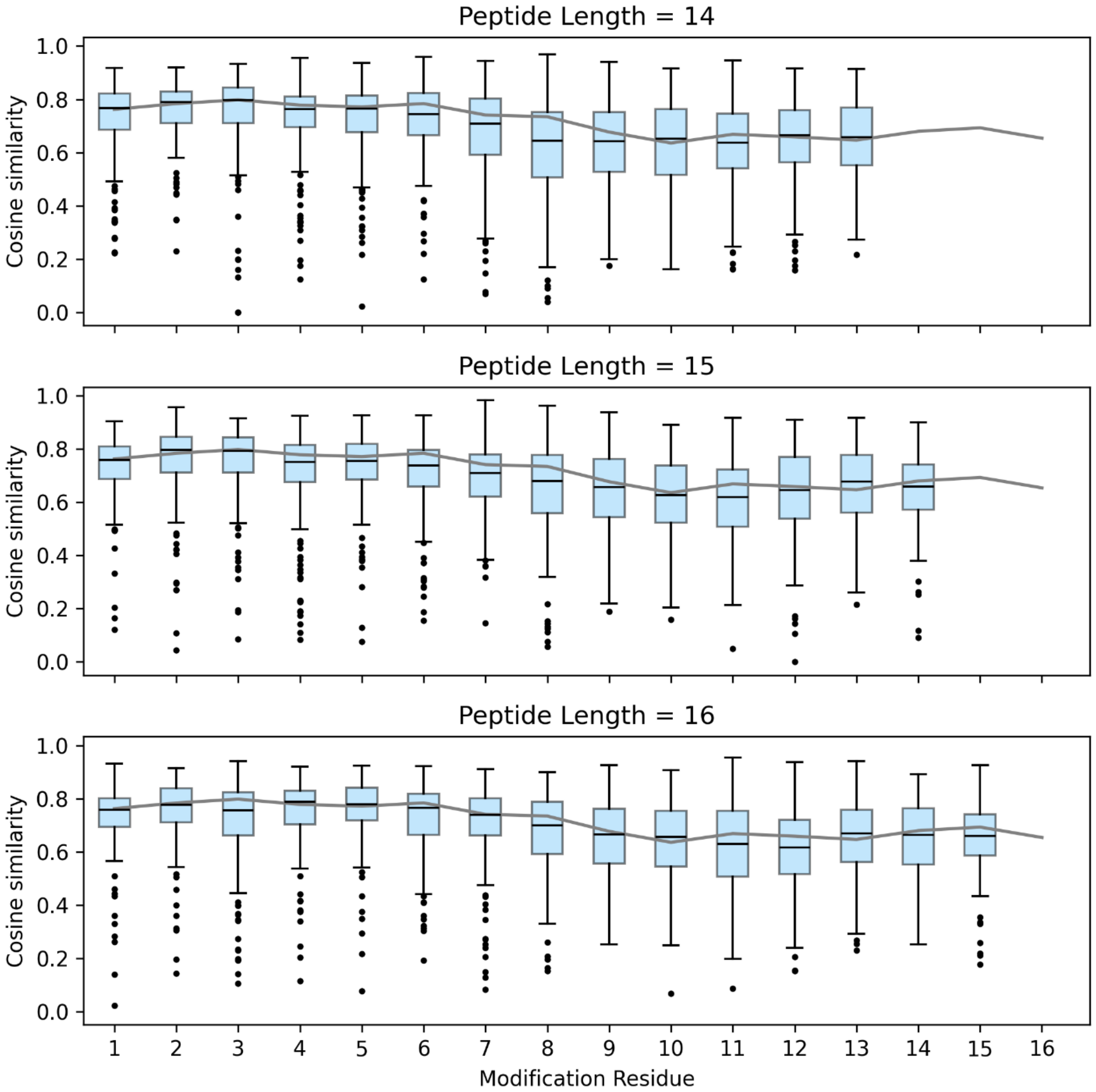


**Figure S3.** Cosine similarity scores for all modified peptides 14, 15, and 16 amino acids long. The solid grey line represents median score values for each modification residue for all modified peptides 17 amino acids long categorized by modification residue. Together, Figures S1-S3 show that as the modification gets closer to the c-terminus of the peptide, similarity score distributions are broadened, and median scores decrease. This trend holds independent of peptide length.

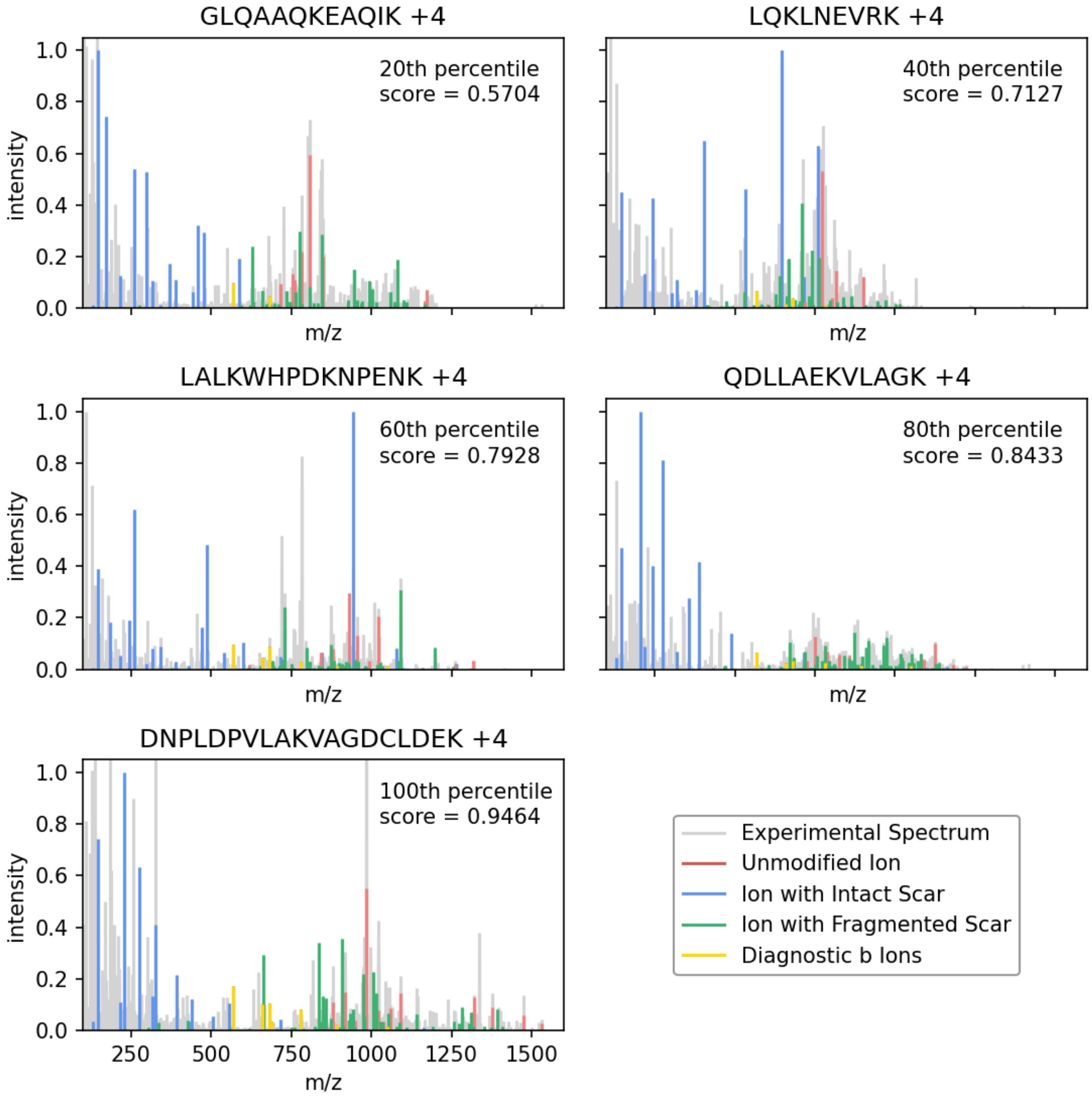


**Figure S4.** Experimental spectra of ubiquitylated LysC peptides with annotated ions corresponding to quintiles based on b-ion scores.

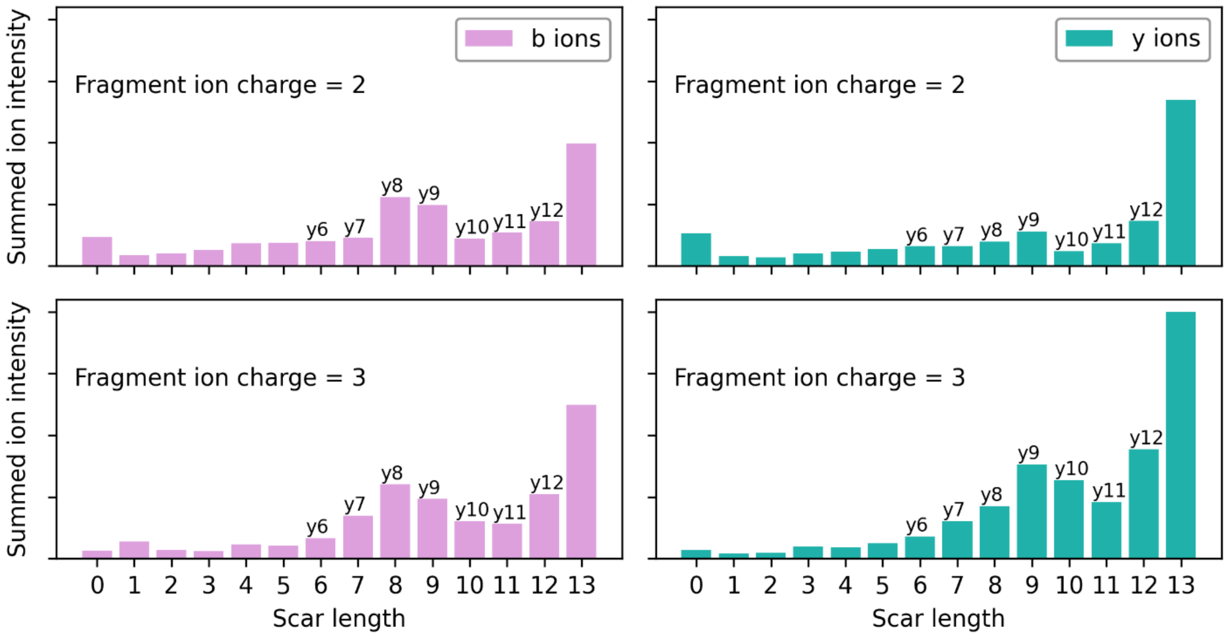


**Figure S5.** Experimental intensities of fragment ions that contain a fragmented Ub scar summed across all experimental spectra. Experimental ions were categorized based on the scar attachment after fragmentation, charge state, and base ion type.
